# UV-C light completely blocks highly contagious Delta SARS-CoV-2 aerosol transmission in hamsters

**DOI:** 10.1101/2022.01.10.475722

**Authors:** Robert J. Fischer, Julia R. Port, Myndi G. Holbrook, Kwe Claude Yinda, Martin Creusen, Jeroen ter Stege, Marc de Samber, Vincent J. Munster

## Abstract

Behavioral and medical control measures are not effective in containing the spread of SARS-CoV-2. Here we report on the effectiveness of a preemptive environmental strategy using UV-C light to prevent airborne transmission of the virus in a hamster model and show that UV-C exposure completely prevents airborne transmission between individuals

## Introduction

The COVID-19 pandemic has officially caused more than 5.4 million deaths worldwide as of December 28, 2021.^1^ Epidemiological and experimental data suggests that the primary mode of transmission of the virus is through airborne particles.^2–4^ Medical countermeasures, such as vaccines and monoclonals antibody therapies were rapidly developed, but have had limited impact on the overall control of the pandemic. While the developed vaccines are highly effective against preventing sever COVID-19 and hospitalization, their transmission-blocking potential on population level appears limited. Currently, 44.65% of the global population are fully vaccinated and an estimated 285 million people have been infected with SARS-CoV-2.^1^ This has drastically changed SARS-CoV-2 immune landscape and likely promoted the emergence of Variants of Concern (VoC) escaping antibody immunity, fueling the current global spikes in infection rates.^5^ These rapid increases in SARS-CoV-2 prevalence prompt crude control measures such as: travel restrictions, large-scale quarantining and “lock downs” of entire populations leading to economic and public health burden.^6^ The inability to control the ongoing SARS-CoV-2 pandemic has put the focus on the development of pathogen agnostic non-medical intervention strategies. These non-medical intervention strategies should ideally be practical, effective under multiple conditions, not depend on the cooperation of individuals, not contribute to virus evolution and prove efficacious for multiple epidemic and pandemic pathogens. One measure that has the potential to decrease the concentration of infectious airborne pathogens in enclosed spaces is ultraviolet (UV) light. Ultraviolet light, in particular UV-C light (wavelengths in the range of 200 nm – 280 nm) has germicidal properties. Several studies have shown that UV-C light can be used to inactivate SARS-CoV-2 on surfaces using a UV-C germicidal lamp.^7–9^ Here we report on the effectiveness of UV-C light in blocking transmission of airborne SARS-CoV-2 in a hamster model.

## Results

To test the ability of UV-C light to prevent infection of naïve hamsters by naturally aspirated aerosols we employed a modified version of an aerosol transmission system described previously.^4^ In this system two cages are separated by a 1250 mm X 73 mm tube resulting in a size exclusion of airborne particles ≥10 μm. The central portion of the tube is quartz enclosed in a HDPE box containing a UV-C light source (Figure 1). The length of the tube inside the box is 66.2 cm and the air traveling from the infected animals to the naïve animals had a residence time of 10.7 seconds in the tube. A 934.5 L/hr airflow, approximately 30 cage air exchanges per hour, is maintained throughout the experiment resulting in a UV-C dose exposure of the pathogen-containing airborne particles of approximately 21.4 mJ/cm^2^.

**Figure 1.**
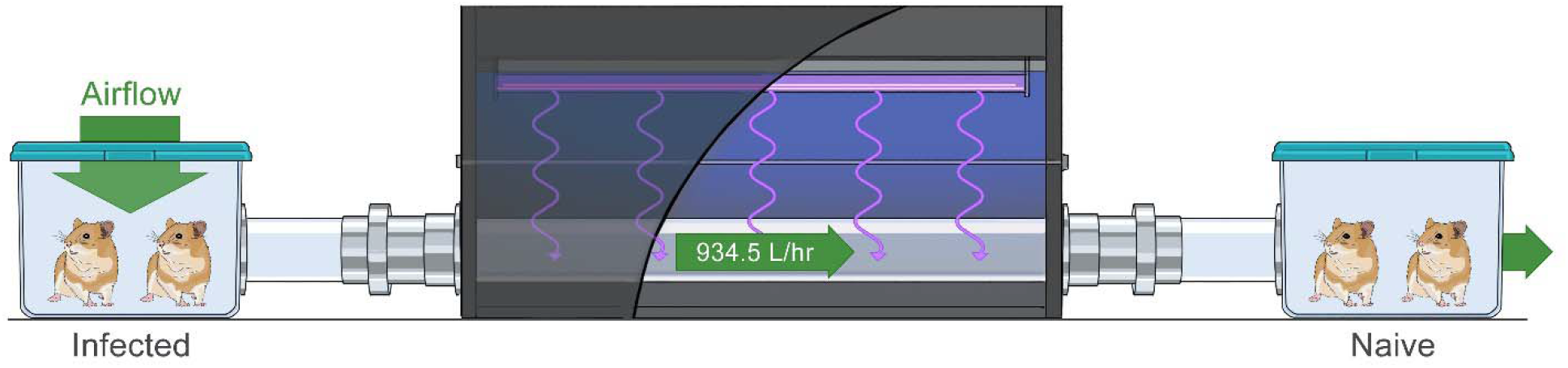
Experimental aerosol transmission with UV-C irradiation setup. Two cages are separated with a 1250 mm X 73 mm i.d. tube. The center portion of the tube is 662 mm of UV transparent quartz surrounded by a HDPE box housing a UV-C light source. Two donor hamsters, infected intranasally with 8 × 10^4^ TCID_50_ SARS-CoV-2 of either lineage A or the Delta variant one day prior to the experiment, were placed in the upstream cage and two naïve sentinel hamsters were placed in the downstream cage with a 934.5 L/hr airflow for 4 hours. The arrow indicates the direction of the airflow.

Briefly, for each trial, 2 hamsters were inoculated intranasally (IN) with 8 × 10^4^ TCID_50_ SARS-CoV-2 strain nCoV-WA1-2020 (EPI_ISL_404895) (prototype lineage A SARS-CoV-2) or hCoV-19/USA/KY-CDC-2-4242084/2021 (EPI_ISL_1823618) (VoC Delta). At 1 day post infection (dpi) 2 infected hamsters were placed in the upstream (donor) cage and 2 naïve hamsters were placed in the downstream (naïve) cage. After a 4-hour exposure the exposed naïve hamsters were moved to individual cages and the donor hamsters were euthanized after an oropharyngeal swab was collected.

To determine whether the naïve exposed sentinel hamsters became infected, oropharyngeal swabs were collected on days 1, 2 and 3 post exposure (DPE) and analyzed for the presence of subgenomic viral RNA (sgRNA, marker for active SARS-CoV-2 replication) and genomic viral RNA (gRNA) by qRT-PCR. The experiment was repeated 4 times for the following conditions: UV-C light treatment, no UV-C light treatment with variant nCoV-WA1-2020 or hCoV-19/USA/KY-CDC-2-4242084/2021 (Delta). When testing under UV-C conditions, the light was turned on 1 hour prior to introducing the animals to the system.

All the animals in the no UV-C treatment groups became infected as early as 1 DPE. gRNA was detected in all animals as early as 1 DPE for both the lineage A and the Delta VOC and continued through DPE 3 (Figure 2, A & C). No gRNA was detected in either of the UV-C groups (Figure 2, A &C). sgRNA was also detected on DPE 1 – 3 in the no UV-C treatment groups, but not in any of the animals in the UV-C groups (Figure 2, B &D). To conclusively demonstrate absence of transmission of SARS-CoV-2 in both UV-C treatment groups the binding antibody titers against the SARS-CoV-2 spike protein (S) were determined on sera obtained at 14 DPE. Both no UV-C light treatment groups had high antibody titers (≥52,000 in all animals, n = 16), but both no UV-C light treatment groups displayed a complete lack of binding antibody titers against SARS-CoV-2 S (<400 in all animals, n = 16).

**Figure 2.**
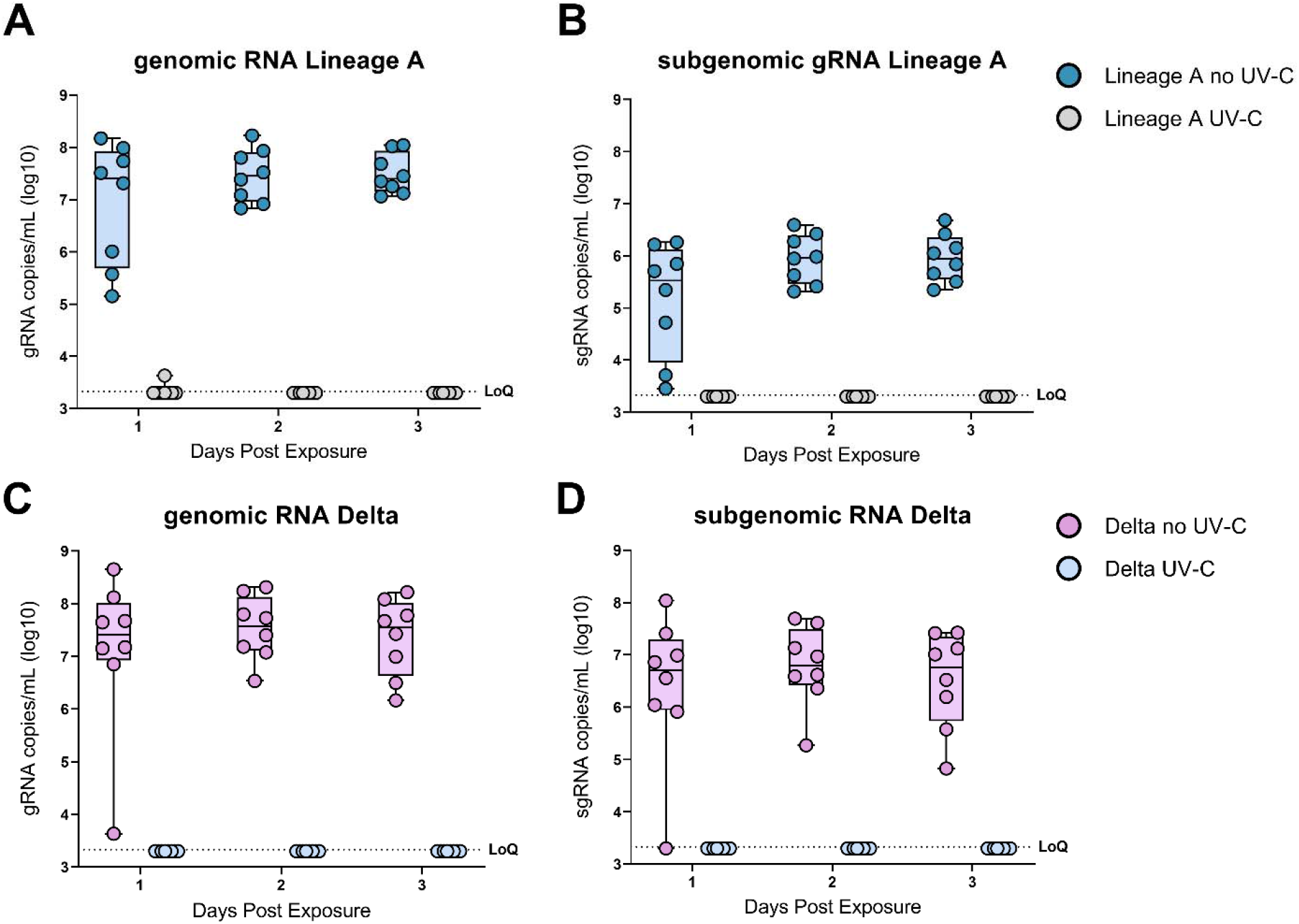
UV-C irradiation blocks SARS-CoV-2 aerosol transmission in hamsters. A & B) Boxplot (minimum to maximum) of genomicRNA and subgenomicRNA Lineage A SARS-CoV-2 RNA in oropharyngeal swabs collected on 1-, 2- and 3-days post exposure. Blue dots represent the no UV-C treatment group (n = 8) and grey dots represent the UV-C treatment group (N=8). C & D) Boxplot (minimum to maximum) of genomicRNA and subgenomicRNA Delta SARS-CoV-2 RNA in oropharyngeal swabs collected on 1-, 2- and 3-days post exposure. Pink dots represent the no UV-C treatment group (n = 8) and light-blue dots represent the represent the UV-C treatment group (N=8). Dotted line = limit of detection.

## Discussion

As the SARS-CoV-2 pandemic approaches its third-year, additional non-medical intervention strategies are urgently needed. Especially in areas and locations where there is a higher risk of SARS-CoV-2 transmission, such as hospitals, COVID-19 testing centers, schools and other indoor areas effective preemptive environmental intervention measures are needed to protect health care workers and people at risk of developing severe COVID19. Non-medical intervention such as social distancing rely on the assumption that small airborne respiration droplets will settle to the ground within about 2 meters from the source. However, true aerosols (< 10 μm) in diameter will remain suspended, floating on air currents for an extended amount of time, can travel more than 2 meters and remain suspended for minutes to hours. In addition, other non-medical countermeasures, such as mask wearing, are highly dependent on compliance and as such have had varying levels of effectiveness across different cultural, political, and religious environments.

Here we have demonstrated that a preemptive environmental intervention measure, using UV-C irradiation, can prevent the aerosol transmission of SARS-CoV-2 between hamsters. This work suggests that UV-C could be used to decrease the concentration of viable air-borne virus in various environments used in conjunction with existing control measures and where other methods are less likely to work. Extensive literature is available for pathogen inactivation, using either bacterial spore inactivation tests, bacteria or respiratory viruses by UV-C treatment.^10,11^ There are several UV-C systems that have been developed and are already being employed.^12,13^ The experiments described here recapitulate a system in which ducted air is treated and returned to the room; the efficiency of this type of system is dependent on the number of room-air exchanges per hour and the ventilation system processes. Another UV-C system that has been employed in areas with a high incidence of tuberculosis (TB) is upper-room ultraviolet germicidal irradiation.^14^ Upper-room ultraviolet germicidal irradiation has the potential to treat up to 24 room air changes per hour equivalents where comfort level ventilation systems handle between 1 and 2 room air exchanges per hour.^15^

Preemptive environmental interventions in public spaces, that are not dependent on the compliance of the at-risk population, would potentially be a highly cost-effective non-medical countermeasure to help control the current pandemic. In addition, given the pathogen agnostic nature of UV-C germicidal irradiation it has the potential to curb airborne transmission of fungal, bacterial, and viral pathogens and even everyday maladies like the common cold.

## Acknowledgements

We would like to thank Neeltje van Doremalen, Jonathan Schulz, Emmie de Wit, Brandi Williamson, Natalie Thornburg, Bin Zhou, Sue Tong, Sujatha Rashid, Ranjan Mukul, Kimberly Stemple, Craig Martens, Kent Barbian, Stacey Ricklefs, Sarah Anzick, Rose Perry and the animal care takers for their assistance during the study. Maarten de Jager, Noud Fleuren, Kevin Clark for their support in the design, construction and calibration of the UV-C apparatus. The following reagent was obtained through BEI Resources, NIAID, NIH: SARS-Related Coronavirus 2, Isolate hCoV-19/England/204820464/2020, NR282 54000, contributed by CDC.

## References

1. WHO. WHO Coronavirus (COVID-19) Dashboard. 12/28/2021. Accessed 12/28/2021. https://covid19.who.int/

2. Kutter JS, de Meulder D, Bestebroer TM, et al. SARS-CoV and SARS-CoV-2 are transmitted through the air between ferrets over more than one meter distance. Nat Commun. Mar 12 2021;12(1)doi:ARTN 1653

3. Zhang RY, Li YX, Zhang ANL, Wang Y, Molina MJ. Identifying airborne transmission as the dominant route for the spread of COVID-19 (vol 117, pg 14857, 2020). P Natl Acad Sci USA. Oct 13 2020;117(41):25942–25943. doi:10.1073/pnas.2018637117

4. Port JR, Yinda CK, Avanzato VA, et al. Increased aerosol transmission for B.1.1.7 (alpha variant) over lineage A variant of SARS-CoV-2. bioRxiv. Jul 26 2021;doi:10.1101/2021.07.26.453518

5. Eguia RT, Crawford KHD, Stevens-Ayers T, et al. A human coronavirus evolves antigenically to escape antibody immunity. Plos Pathogens. Apr 2021;17(4)e1009453. doi:10.1371/journal.ppat.1009453

6. Tisdell CA. Economic, social and political issues raised by the COVID-19 pandemic. Economic Analysis and Policy. Dec 2020;68:17–28. doi:10.1016/j.eap.2020.08.002

7. Fischer R, Morris D, van Doremalen N, et al. Effectiveness of N95 Respirator Decontamination and Reuse against SARS-CoV-2 Virus. Emerging Infectious Disease journal. 2020;26(9):2253. doi:10.3201/eid2609.201524

8. Storm N, McKay LGA, Downs SN, et al. Rapid and complete inactivation of SARS-CoV-2 by ultraviolet-C irradiation. Sci Rep-Uk. Dec 30 2020;10(1)doi:ARTN 22421 10.1038/s41598-020-79600-8

9. Ruetalo N, Businger R, Schindler M. Rapid, dose-dependent and efficient inactivation of surface dried SARS-CoV-2 by 254 nm UV-C irradiation. Eurosurveillance. Oct 21 2021;26(42)doi:Artn 2001718 10.2807/1560-7917.Es.2021.26.42.2001718

10. Browne K. Brought to Light: How Ultraviolet Disinfection Can Prevent the Nosocomial Transmission of COVID-19 and Other Infectious Diseases. Applied Microbiology. 2021;1(3):537–556.

11. Xu P, Peccia J, Fabian P, et al. Efficacy of ultraviolet germicidal irradiation of upper-room air in inactivating airborne bacterial spores and mycobacteria in full-scale studies. Atmospheric Environment. Jan 2003;37(3):405–419. doi:Pii S1352-2310(02)00825-7 Doi 10.1016/S1352-2310(02)00825-7

12. Ethington T, Newsome S, Waugh J, Lee LD. Cleaning the air with ultraviolet germicidal irradiation lessened contact infections in a long-term acute care hospital. Am J Infect Control. May 2018;46(5):482–486. doi:10.1016/j.ajic.2017.11.008

13. Qiao YC, Yang M, Marabella IA, et al. Greater than 3-Log Reduction in Viable Coronavirus Aerosol Concentration in Ducted Ultraviolet-C (UV-C) Systems. Environmental Science & Technology. Apr 6 2021;55(7):4174–4182. doi:10.1021/acs.est.0c05763

14. First M, Rudnick SN, Banahan KF, Vincent RL, Brickner PW. Fundamental factors affecting upper-room ultraviolet germicidal irradiation - part I. Experimental. J Occup Environ Hyg. May 2007;4(5):321–31. doi:10.1080/15459620701271693

15. Nardell EA. Air Disinfection for Airborne Infection Control with a Focus on COVID-19: Why Germicidal UV is Essential(dagger). Photochem Photobiol. May 2021;97(3):493–497. doi:10.1111/php.13421

