## supplemental data and material and methods for "UV-C light completely blocks highly contagious Delta SARS-CoV-2 aerosol transmission in hamsters"

**Supplemental Online Materials**

**Background:**

*Mechanism of UV-C inactivation*

UV-C is effective in inactivating microorganisms such as fungi, bacteria and viruses including SARS-CoV-2^1^. UV-C light is primarily generated by highly efficient and well-established low-pressure (LP) mercury discharge lamps, that emit a 254 nm peak wavelength, which is close to the optimal UVGI wavelength of 260-265 nm^2^. The mechanism of UV-C deactivation of viruses and bacteria is first a photochemical cross-linking reaction of nucleic acids, which prevents transcription and replication of RNA/DNA in host cells^3^. A second photochemical reaction targets proteins, leading to protein molecular modifications which result in e.g., loss of host cell recognition of the virus and damage to membranes and envelopes^2^.

**Material and Methods:**

*Ethics Statement*

All animal work was performed in an AAALAC International accredited facility, under animal study protocol (ASP) # 2021-015 as approved by the RML Animal Care and Use Committee (ACUC) in accordance with guidelines set forth in the Guide for the Care and Use of Laboratory Animals 8th edition, the Animal Welfare Act, United States Department of Agriculture and the United States Public Health Service Policy on the Humane Care and Use of Laboratory Animals.

The experiments were conducted in a BSL4 facility and the samples were removed following approved SOPs.

*Aerosol transmission apparatus*

To determine if UV light is capable of arresting aerosol transmission between hamsters, we modified the aerosol transmission system described previously by Port et al^4^. Two rodent boxes (Lab Products Inc.), one designated the donor box and the other the naïve box, were connected by a 1250 mm long by 73 mm inside diameter, UV-C transparent, quartz tube containing stainless steel mesh at each end to prevent the hamsters from traversing the tube. The tube passed through a 662 mm long HDPE box that housed the UV-C light source and shielded the surrounding area and animals from incident UV-C light. UV-C light was generated with two 58.7 cm Philips TUV F17T8 mercury lamps preaged to provide a stable UV-C light output. The center of the quartz tube was positioned at 205 mm from the UV-C source, 59 mm from the floor of the box and 131 mm from the walls of the box.

A directional air flow, from the donor box to the naïve box was created using a vacuum pump connected to the naïve box and controlled with a float type meter/valve combination (King Industries). Air entered the system through the filtered lid of the donor box while the lid of the naïve box was fitted with an air impermeable film to ensure the air flowed from the donor box to the naïve box.

Air was drawn through the system at 934.5 L/hr. representing approximately 30 air exchanges/hr.

The velocity of the air as it moves through the connecting tube was calculated by the following:

eq. 1

$$\frac{V_{m}}{{A'}_{tube}}=v$$

where *V_m_* is the volume of air traveling through the connecting tube in cm^3^/min and *A′_tube_* represents the cross section of the connecting tube in cm^2^. The amount of time (*t*) needed for a front of air to traverse the tube is then calculated using:

eq. 2

$$\frac{L}{v}=t$$

where *L* is the length of the tube in cm and *v* is the velocity in cm/min. calculated in eq.1.

The incidence of UV-C light at 254 nm was measured at 4 points along the length of the tube at 20 cm intervals starting 1 cm from where the tube exits the UV-C containment box with a UV-C meter type X1-1-UV-3725 measurement system, comprising a X1-5 Optometer with UV-3725-5 Detector head, calibrated for narrow band sources such as LP mercury lamps. After a 1-hour lamp warmup, irradiance measurements (mW/cm^2^) were taken in triplicate with the sensor facing 1) the UV-C light source (top), 2) away from the UV-C light source (bottom) 3) the right side of the box and 4) the left side of the box (Supplemental Table 1). These irradiance dose measurements were used to calculate the total UV-C incidence dose (mJ/cm^2^) along the length of the tube by multiplying irradiance dose by exposure time.

*Total UV incidence and UV-C dose*

To calculate the UV-C dose that pathogens travelling through the tube experience an approximation was employed by using only the upward-registered irradiance values at the 4 measurement points. This simplification was used as these values are the main contributors. The resulting overall average dose is hence a minimal value. The upward facing UV-C irradiances at the 4 measurement points were plotted and a curve fitted to the points. The average UV-C irradiance was calculated by taking the integral from 0 to 66.2 cm. (Supplemental Figure 1)

*Cells and virus*

SARS-CoV-2 variant SARS-CoV-2 strain nCoV-WA1-2020 (Lineage A, EPI_ISL_404895) was obtained from Nathalie Thornburg at CDC. SARS-CoV-2 hCoV-19/USA/KY-CDC-2-4242084/2021 (Delta, B.1.617.2, EPI_ISL_1823618) was obtained from BEI resources. All virus stocks were sequenced, no SNPs compared to the original patient sample sequence were detected. Virus propagation was performed in VeroE6 cells in DMEM supplemented with 2% fetal bovine serum, 1 mM L-glutamine, 50 U/ml penicillin and 50 μg/ml streptomycin (DMEM2). VeroE6 cells were maintained in DMEM supplemented with 10% fetal bovine serum, 1 mM L-glutamine, 50 U/ml penicillin, and 50 μg/ml streptomycin. No mycoplasma was detected in cells or virus stocks.

*Hamster to hamster transmission*

Male and female Syrian golden hamsters 6 to 7 weeks old were used in these experiments. Sixteen hamsters were used for each experimental group. Hamster to hamster aerosol transmission was evaluated using 2 donor hamsters and 2 naïve hamsters for each of 4 replicates for every virus/condition. One day prior to the experimental exposure donor hamsters were inoculated intranasally with 8 X 10^4^ TCID_50_ SARS-CoV-2 Lineage A or the Delta variant. In the trials where UV-C light was being used the UV-C lamps were turned on one hour prior to the exposure. To start the experiment the naïve animals were placed into their box, then the box was connected to the system, the same was done for the donor animals. To start the experiment the pump was turned on to produce an air flow from the donor cage to the naïve cage. The animals remained in the transmission system for 4 hours, then the naïve animals were singly housed in standard caging. The donor animals were swabbed upon the completion of the 4-hour exposure, the naïve exposed animals were swabbed for 3 consecutive days starting on day 1 post exposure. On day 14 blood was drawn from the exposed naïve animals for serology and the experiment was concluded.

*RNA extraction and quantitative reverse-transcription polymerase chain reaction*

Oropharyngeal swabs were collected from anesthetized hamsters and placed in 1 mL of cell culture medium. RNA was extracted using a Qiagen viral RNA 96-well format kit on a Qiacube robot according to the manufacturer’s instructions and following high containment laboratory protocols. qRT-PCR was carried out using Taqman fast-virus 1-step master mix with previously described E_Sarbeco or primers and probe (E gene) or sgLeadSARS2-F forward primer with E_Sarbeco reverse primer and probe (subgenomicE gene), on a Quant studio 3.^5^

*ELISA assay:*

ELISA assays were performed as previously described.^5^ In brief, maxisorp plates (Nunc) were coated with 50 ng spike protein (Sinobiological SARS-CoV-2 (2019-nCoV) spike S1 + S2 ECD-HIS recombinant protein (catalog #40589-V08B1)) per well. Plates were incubated overnight at 4°C. Plates were blocked with casein in phosphate buffered saline (PBS) (ThermoFisher) for 1 hours at room temperature (RT). Hamster serum collected on day 14 post exposure was diluted 1:400 in in casein in PBS and incubated for 1 hours at RT. Secondary goat anti-hamster IgG Fc (horseradish peroxidase (HRP)-conjugated, Abcam) spike-specific antibodies were used for detection and visualized with KPL TMB 2-component peroxidase substrate kit (SeraCare, 5120-0047). The reaction was stopped with KPL stop solution (Seracare) and plates were read at 450 nm. The threshold for positivity was calculated as the average plus 3 x the standard deviation of negative control hamster sera.

**References**

1.	Biasin M, Bianco A, Pareschi G, et al. UV-C irradiation is highly effective in inactivating SARS-CoV-2 replication. *Sci Rep-Uk*. Mar 18 2021;11(1)doi:ARTN 6260

2.	Kowalski W. Ultraviolet Germicidal Irradiation Handbook. *Ultraviolet Germicidal Irradiation Handbook*. 2009:1-501. doi:10.1007/978-3-642-01999-9

3.	Cutler TD, Zimmerman JJ. Ultraviolet irradiation and the mechanisms underlying its inactivation of infectious agents. *Anim Health Res Rev*. Jun 2011;12(1):15-23. doi:10.1017/S1466252311000016

4.	Port JR, Yinda CK, Avanzato VA, et al. Increased aerosol transmission for B.1.1.7 (alpha variant) over lineage A variant of SARS-CoV-2. *bioRxiv*. Jul 26 2021;doi:10.1101/2021.07.26.453518

5.	Fischer RJ, van Doremalen N, Adney DR, et al. ChAdOx1 nCoV-19 (AZD1222) protects Syrian hamsters against SARS-CoV-2 B.1.351 and B.1.1.7. *Nat Commun*. 2021/10/07 2021;12(1):5868. doi:10.1038/s41467-021-26178-y

**eTable 1.** UV-C light incidence at 254 nm at 4 points along the length of the quartz tube at 20 cm intervals starting 1 cm from the UV-C containment box exit. Measurements were made with a UV-C meter type X1-1-UV-3725 measurement system, comprised of a X1-5 Optometer with UV-3725-5 Detector head, calibrated for narrow band sources such as LP mercury lamps. After a 1-hour lamp warmup, irradiance measurements (mW/cm^2^) were taken in triplicate with the sensor facing 1) the UV-C light source (top), 2) away from the UV-C light source (bottom) 3) the right side of the box and 4) the left side of the box (Supplemental Table 1). These irradiance dose measurements were used to calculate the total UV-C incidence dose (mJ/cm^2^) along the length of the tube by multiplying irradiance dose by exposure time.

**eFigure 1.** The average of three upward facing UV-C irradiance measurements (Top, eTable 1) were plotted and a second order polynomial curve was fitted to the points. The average UV-C irradiance was calculated by taking the integral from 0 to 66.2 cm.

**eTable 1.**

| Distance from quartz tube exit | Top | Left | Bottom | Right |
| --- | --- | --- | --- | --- |
| 1 cm | 0.994 | 0.114 | 0.018 | 0.100 |
| 21 cm | 2.437 | 0.245 | 0.042 | 0.240 |
| 41 cm | 2.555 | 0.282 | 0.073 | 0.315 |
| 61 cm | 1.345 | 0.207 | 0.093 | 0.250 |

**eFigure 1.**

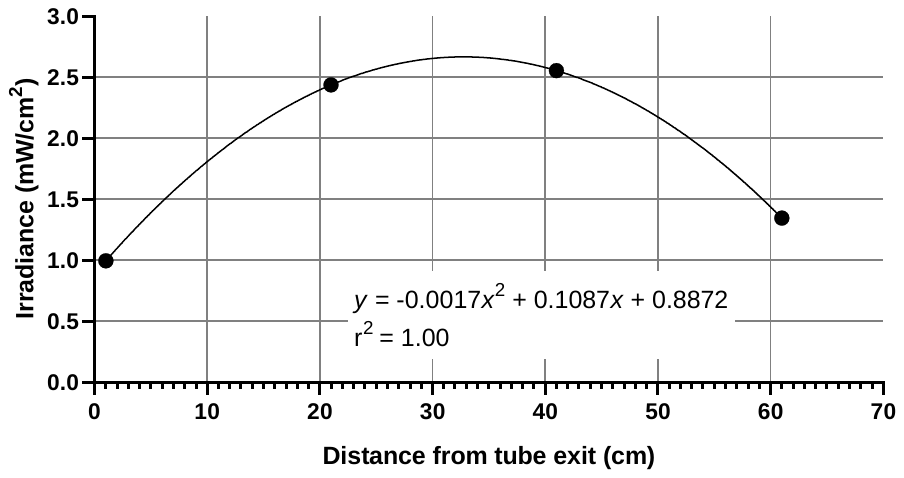
